# Aberrant Functional Connectivity between Reward and Inhibitory Control Networks in Pre-Adolescent Binge Eating Disorder

**DOI:** 10.1101/2021.10.06.463386

**Authors:** Stuart B. Murray, Celina Alba, Christina J. Duval, Jason M. Nagata, Ryan P. Cabeen, Darrin J. Lee, Arthur W. Toga, Steven J. Siegel, Kay Jann

## Abstract

**Background:** Behavioral features of binge eating disorder (BED) suggest abnormalities in reward and inhibitory control. Studies of adult populations suggest functional abnormalities in reward and inhibitory control networks. Despite behavioral markers often developing in children, the neurobiology of pediatric BED remains unstudied.

**Methods:** 58 pre-adolescent children (aged 9-10-years) with BED and 66 age, BMI and developmentally-matched control children were extracted from the 3.0 baseline (Year 0) release of the Adolescent Brain Cognitive Development (ABCD) Study. We investigated group differences in resting-state functional MRI (rs-fMRI) functional connectivity (FC) within and between reward and inhibitory control networks. A seed-based approach was employed to assess nodes in the reward (orbitofrontal cortex, nucleus accumbens, amygdala) and inhibitory control (dorsolateral prefrontal cortex, anterior cingulate cortex) networks via hypothesis-driven seed-to-seed analyses, and secondary seed-to-voxel analyses.

**Results:** Our findings revealed reduced FC between the dlPFC and amygdala, and between the anterior cingulate cortex and orbitofrontal cortex in pre-adolescent children with BED, relative to age, gender, BMI and developmentally matched controls. These findings indicating aberrant connectivity between nodes of inhibitory control and reward networks were corroborated by the whole-brain FC analyses.

**Conclusions:** Early-onset BED may be characterized by diffuse abnormalities in the functional synergy *between* reward and cognitive control networks, without perturbations *within* reward and inhibitory control networks, respectively. The decreased capacity to regulate a reward-driven pursuit of hedonic foods, which is characteristic of BED, may in part, rest on this dysconnectivity between reward and inhibitory control networks.

---

Eating disorders (ED) affect more than 9% of the US population in their lifetime, which equates to more than 28 million Americans (1). This constellation of pernicious and burdensome psychiatric disorders typically run a chronic and relapsing illness course and result in more than 10,000 deaths per year in the United States (1). The most prevalent of all ED phenotypes is binge eating disorder (BED), which affects up to 3-5% of the US population (1). BED is characterized by (i) frequent consumption of an objectively large amount of food in a discrete time period, (ii) a subjective and self-reported loss of control, and (iii) an absence of compensatory behaviors to offset caloric intake (2). Importantly, BED portends an array of deleterious medical and psychiatric sequalae, including weight gain, metabolic syndrome, diabetes, hypertension, dyslipidemia, abnormal cardiac function, and elevated suicidality (3). With specific regards to weight, more than 75% of those with BED are overweight or obese (3), which in itself ranks among the leading causes of preventable death on a global scale. With treatment options for BED remaining limited (4), and predominantly behaviorally focused (4), the need to identify and target the mechanisms underpinning BED psychopathology is critical.

Interestingly, whereas dietary restraint typically *precedes* binge eating behaviors in other ED presentations (5), this is typically not the case in BED, where a developmental emergence of binge episodes precedes attempts at dieting (5). Indeed, low pre-meal levels of the pro-appetitive hormone ghrelin level further suggests an absence of homeostatic antecedents to binge episodes in those with BED (6). Instead, the heightened drive towards high-fat and high-sugar foods during binge episodes (7) when not calorically restricted suggests altered reward sensitivity and impulsivity in BED. Hedonic eating - the intense pleasure derived from eating highly palatable food, even when not hungry or calorically deprived, is elevated among those with BED (8), and prospectively predicts the frequency and intensity of food cravings and binge episodes (9). In concert, elevated food-related impulsivity (10) and difficulty diverting attention away from food-related cues (10) is evident among those with BED (10), and is intricately linked to the inability to abstain from food consumption (i.e., loss of control eating) which characterizes BED. Broad deficits in behavioral inhibition have been noted, although this is particularly evident with respect to food stimuli (10). Interestingly, elevated hedonic eating appears to drive greater palatable (as opposed to bland) food consumption when low inhibitory control is present (11), suggesting a critical intersectionality of altered reward and inhibitory control-related processes in BED.

Mechanistically, the neural circuitry underpinning reward and inhibitory control-related processes is well defined. The anticipation and receipt of reward and the initiation of goal-directed behavior involve a well-defined neural circuit comprising the anterior cingulate cortex, amygdala, ventral tegmental area (VTA), bed nucleus of the stria terminalis (BNST), orbitofrontal cortex (OFC), ventral striatum (VS)/nucleus accumbens (NAcc), and the prefrontal cortex (PFC) (12). Upon receipt of hedonic cues, the experience of pleasure appears to converge in the ventral striatum (VS) and orbitofrontal cortex (OFC), where μ-opioid and endocannabinoid receptors mediate the hedonic perception of reward. Inhibitory control processes that suppress the dominant or habitual response to cue and goal-directed behavior are associated with activity in the prefrontal cortex: dorsolateral prefrontal cortex (dlPFC), inferior frontal cortex (IFC) and anterior cingulate cortex (ACC). The prefrontal cortex receives input from the VTA, thalamus and amygdala and in turn modulates reactivity of those areas through top-down control over the serotonergic and dopaminergic neurotransmitter systems (13).

To date, relatively few neuroimaging studies have assessed the neural circuits involved in BED psychopathology (14), and those that exist have predominantly assessed adults. Existing studies of those engaging in binge eating episodes have revealed no consistent alterations in gray matter morphometry (15), although arterial spin labeling studies suggest that women with binge type eating disorder presentations (BED; BN) demonstrate increased regional cerebral blood flow in both reward and inhibitory control regions (16). Functional MRI (fMRI) studies have illustrated elevated orbitofrontal cortex (OFC) and ventral striatum (VS) activity during food cue presentation tasks (17), which predicts clinical BED symptom severity (18). In contrast, however, non-food-related reward processing tasks illustrate *attenuated* striatal activity during reward anticipation (19), which is consistent with the blunted anticipatory processing observed in other disorders characterized by impulsivity (20). During inhibitory control tasks, BED is characterized by reduced activity in nodes involved in inhibitory control, such as the prefrontal cortex (PFC) and inferior frontal gyrus (IFG), relative to obese patients and healthy controls (21). Moreover, PFC and IFG hypoactivity is directly related to impaired dietary restraint in those with BED (21), suggesting that the diminished recruitment of impulse-control circuitry is related to an impaired ability to limit food intake.

Few studies to date have assessed functional connectivity in BED. One recent study of adults with BED noted hypoconnectivity between striatal regions which regulate reward processing, and prefrontal regions involved in cognitive and executive control (22). Moreover, this hypoconnectivity was associated with impaired performance on a compulsive reward seeking task, and with increased binge frequency (22), suggesting that a disruption in the functional architecture of reward networks may drive BED psychopathology. In another study of adults with obesity, some of whom had BED, frontostriatal hypoconnectivity was also evident, relative to non-obese controls (23). However, while providing critical preliminary insights into the functional architecture of BED, these studies have been limited by relatively small sample sizes, and an exclusive focus on adult presentations of BED. Despite clinical markers of BED being evident in pre-adolescent childhood (i.e., 6-12 years of age) (24), and early onset BED being associated with greater psychiatric and medical morbidity (25), no study to date has assessed the neural correlates of BED in children.

The present study aimed to leverage the Adolescent Brain Cognitive Development (ABCD) Study (26) to undertake the first known assessment of resting state functional connectivity in pre-adolescent children with BED. In particular, we aimed to assess functional connectivity within and between nodes of the reward and inhibitory control networks in pre-adolescent children with BED, relative to age- and developmentally matched control pre-adolescent children. Owing to the centrality of altered reward and cognitive control circuit activity in presentations of BED, alongside recently demonstrated fronto-striatal dysconnectivity, it was hypothesized that reduced resting state functional connectivity would be evident in children with BED, within and between both reward and inhibitory control-related networks, respectively.

## Methods

### Study Sample

The ABCD Study is a large, diverse, and prospective cohort study of brain development and health throughout adolescence. We analyzed data from the ABCD 3.0 release from baseline (Year 0), which consisted of 11,875 pre-adolescent children aged 9-10 years collected in 2016-2018, recruited from 21 sites around the U.S. Further details of the study sample, recruitment process, exclusion criteria, procedures, and measures have been previously reported (27). Centralized institutional review board (IRB) approval was obtained from the University of California, San Diego. Study sites obtained approval from their local IRBs. Caregivers provided written informed consent and each child provided written assent.

### Measures

#### Diagnostic Interview

Children and parents/caregivers completed the eating disorder module of the Kiddie Schedule for Affective Disorders and Schizophrenia (KSADS-5) (28), assessing frequency, duration, and associated distress of their child’s eating behavior. We used parent/caregiver reports of child behavior due to challenges presented for young children in rating complex cognitive constructs such as loss of control. This approach has been validated based on evidence noting that parents are particularly important reporters for these behaviors in this age range (29). Diagnoses were made according to DSM-5 criteria for BED (2).

#### Pubertal Development Scale (PDS)

Owing to the narrow age range of this sample (9-10 years), and the differential rates of neurodevelopmental maturation among pre-adolescent children of the same chronological age, a self-reported and parent-reported measure of pubertal status (30), frequently used as a measure of developmental maturation when studying brain function and structure (31), was used in the present study to control for differential rates of maturation.

#### Body Mass Index

BMI was calculated based on the average of two-to-three measured heights and weights (BMI = weight/height^2^) by research staff.

### MRI Data Acquisition and Preprocessing

Structural and resting state fMRI data used in this study were from the minimally preprocessed imaging data in the ABCD study as provided on NIH Data Archive (Release 3.0). Structural T1 weighted images used for normalization and atlas parcellation were collected on Siemens Prisma (TR/TE= 2500/2.88ms, 176 slices, 256×256 matrix size, voxel size 1×1×1mm^3^, FA= 8^°^), Philips Achieva (TR/TE= 6.31/2.9ms, 225 slices, 256×240 matrix size, voxel size 1×1×1mm^3^, FA= 8^°^) and GE MR750 (TR/TE= 2500/2ms, 208 slices, 256×256 matrix size, voxel size 1×1×1mm^3^, FA= 8^°^) scanner platforms. Resting-state functional MRI (rs-fMRI) data was acquired with the same imaging parameters across the three scanner platforms: TR/TE= 800/30ms, 60 slices, 216×216 matrix size, voxel size 2.4×2.4×2.4mm^3^, FA= 52^°^, MB= 6, Volumes= 383.

MRI data was analyses in ConnToolbox (32). Preprocessing included motion alignment and bandpass filtering between 0.01Hz and 0.1Hz. To account for physiological noise, we corrected for signal from the white matter and cerebrospinal fluid from the functional data using aCompCor (33). Furthermore, data was motion scrubbed which is particularly salient in pediatric populations (34). Noise corrected fMRI images were then coregistered to anatomical T1 weighted images, normalized into MNI standard space and split into ROIs based on the Harvard-Oxford atlas.

### MRI Data Analysis

Seed-based functional connectivity analysis for selected ROIs representing key nodes of the inhibitory control and reward networks (i.e. orbito-frontal cortex, anterior cingulate cortex, nucleus accumbens, dorsolateral prefrontal cortex, and amygdala, Fig1A) provided subject specific seed-to-seed connectivity matrices. Second level analysis to test for group differences between the BED and control (CONT) groups included the subject specific connectivity matrices as dependent variable, group as independent variable and covariates for pubertal status (PDS), gender and BMI. Significance was set at p<0.05 with correction for 36 (all possible connections between 9 ROIs) multiple comparisons using Benjamini-Hochberg FDR correction at q<0.05. Additional seed-to-voxel functional connectivity analyses were performed to reveal whole-brain patterns of connectivity between ROIs within the inhibitory control and reward networks with brain areas in the rest of the brain. Second-level analysis of the seed-to-voxel models mirrored the seed-to-seed model as it evaluated the group differences between the BED and CONT groups, controlling for pubertal status (PDS), gender and BMI as covariates. Significance was set at p<0.05 with correction for multiple comparisons using false discovery rate (FDR q<0.05).

**Figure 1:**
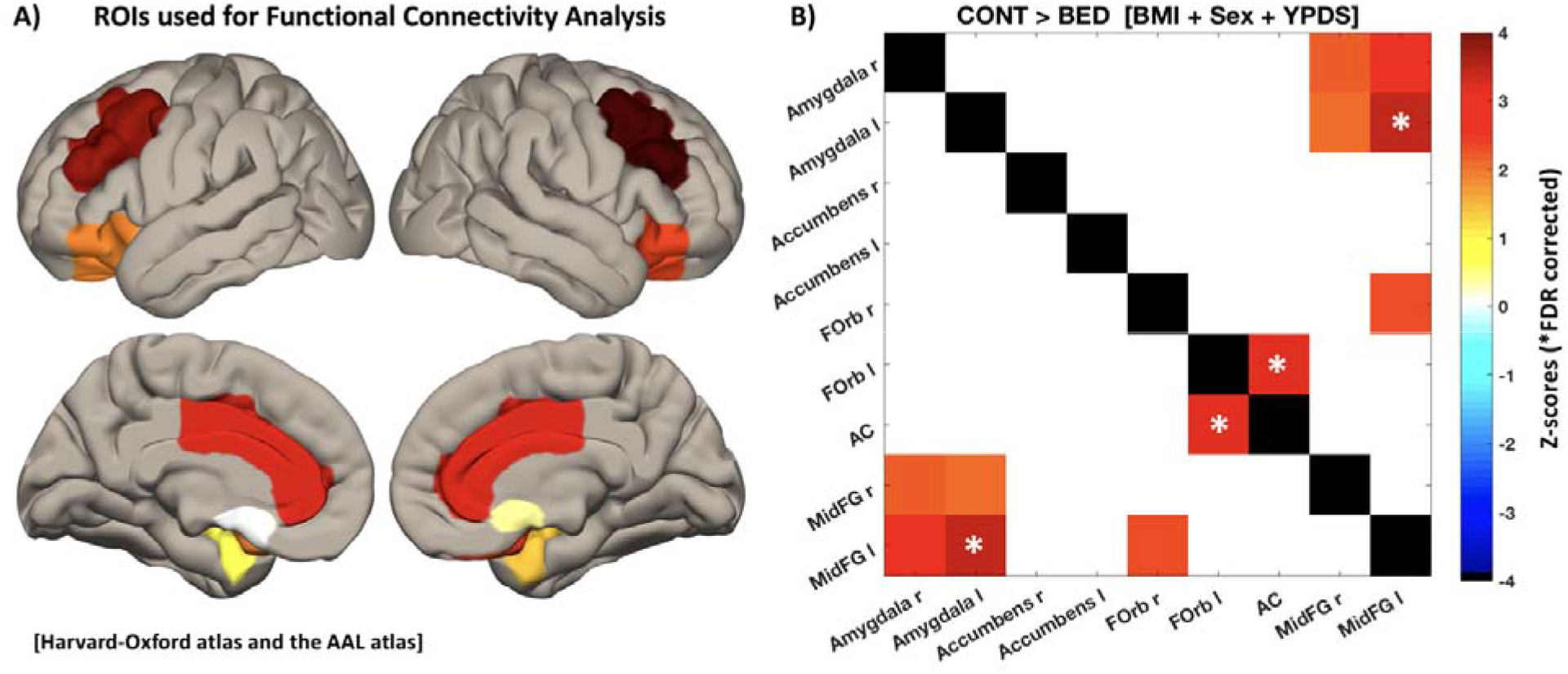
Seed-to-seed rs-fMRI analyses comparing pre-adolescent children with binge eating disorder and control pre-adolescent children, while adjusting for BMI and pubertal development. Seed regions included bilateral amygdala, nucleus accumbens, orbitofrontal cortex (FOrb), dorsolateral prefrontal cortex (MidFG), and the anterior cingulate (AC). Darker red coloring indicates a more positive z-score, and darker blue indicates a more negative z-score (of which there were no significant effects).

## Results

### Sample Demographics

We identified 58 pre-adolescent children (28 females; 30 males) with BED diagnoses based on DMS-5 criteria in the baseline visit of the ABCD study. We also selected 68 healthy control pre-adolescent children (36 females; 32 males) that matched the BED cohort in age (p=0.1197), gender (p=0.6016), BMI (p=0.1617) and PDS (p=0.4854). Table 1 illustrates a detailed characterization of the two cohorts. Among the overall sample, approximately 58% identified as White, 16% as Black, 14% as Mixed Race, 1% as Asian, and 11% as ‘other’. Additionally, 24% of the sample identified as Hispanic.

**Table 1:**
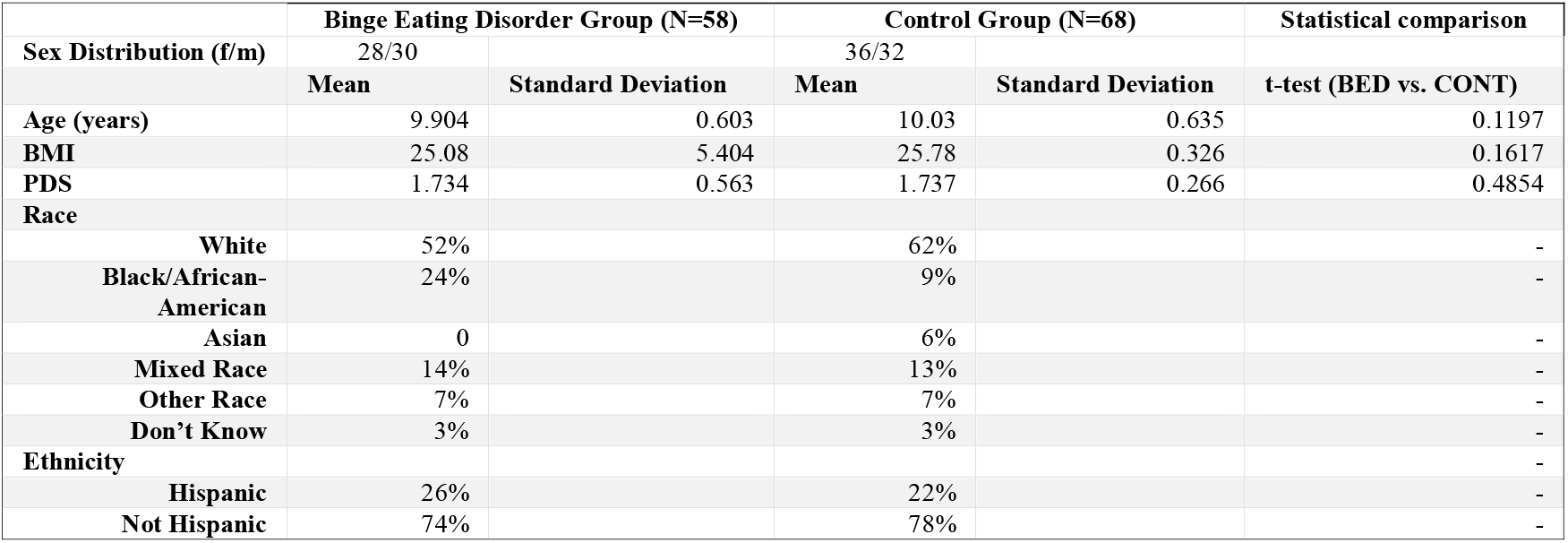
An overview of sample characteristics from the Adolescent Brain Cognitive Development Study, delineated by group.

### Seed-to-Seed connectivity

Statistical analysis between the BED cohort (BED) and control group cohort (CONT) for specific seed-to-seed connectivity revealed significantly lower connectivity between areas of the inhibitory control and reward network. Specifically, left orbitofrontal cortex (OFC) to anterior cingulate cortex (ACC), bilateral dorsolateral prefrontal cortex (DLPFC) to bilateral amygdala (Amyg) and right OFC to left DLPFC (Fig1B). However, only two of these findings survived correction for multiple comparison: AC to L-OFC and L-DLPFC to R-OFC (indicated by asterisk in Fig1B). Notably, all findings indicate differences in the between network connectivity as we found no significant differences for connections between ROIs within the reward or cognitive control network, respectively.

### Seed-to-Voxel connectivity

Statistical analysis of the connectivity from each bilateral pair of ROIs to all other voxels in the brain resulted in connectivity patterns indicating widespread hypo-connectivity in the BED group as compared to the CONT group (Fig2). For the amygdala seeds we saw hypo connectivity in the BED group to DLPFC, PCC, superior frontal gyrus (SFG) and temporal lobe and a small cluster of hyper-connectivity to the temporal pole. The OFC showed reduced connectivity to PCC, ACC, DLPFC and ventromedial prefrontal cortex (VMPFC). The ACC displayed hypo connectivity to OFC, SFG, parahippocampal gyrus and inferior temporal gyrus. Finally, the dlPFC exhibited reduced connectivity to OFC, middle temporal gyrus, SFG and inferior frontal gyrus (IFG). We did not find any significant group differences for the Nacc seed.

**Figure 2:**
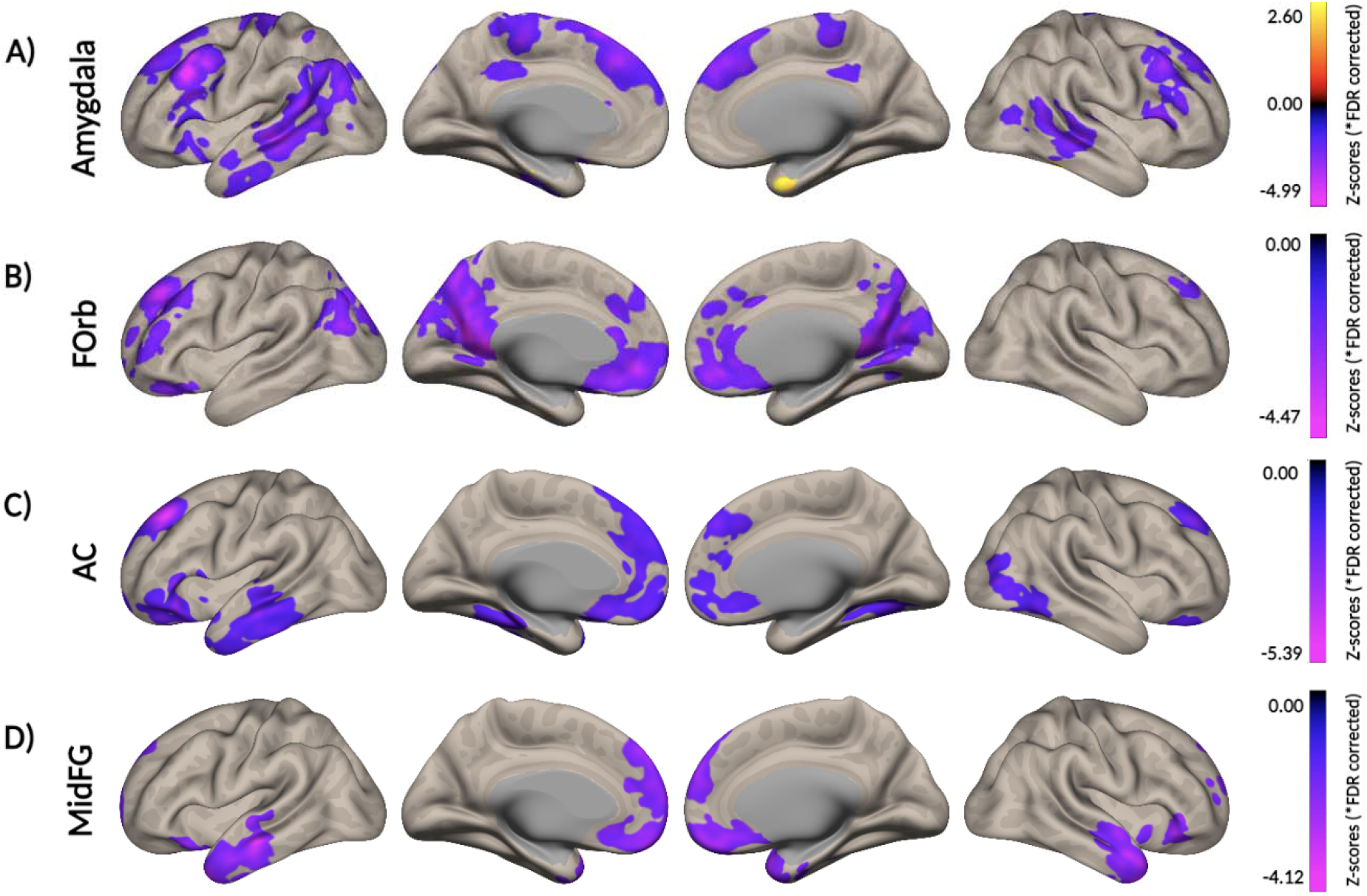
Seed-to-voxel analyses comparing pre-adolescent children with binge eating disorder and control pre-adolescent children, while adjusting for BMI and pubertal development. Seed regions included (A) the amygdala, (B) the orbitofrontal cortex (FOrb), (C) the anterior cingulate (AC), and (D) the dorsolateral prefrontal cortex (MidFG). Hot colors in red, yellow, and white indicate a more positive z-score, and cooler colors in blue, purple, and pink indicate a more negative z-score. Gray brain areas showed no significant difference between groups.

## Discussion

This study reported the first known examination of the functional organization and interaction within and between reward and cognitive control networks in pre-adolescent children with BED, using hypothesis-driven seed-based rsFC analyses. With theoretical and task-evoked neuroimaging evidence suggesting that BED may be characterized by abnormal neural activity in these networks (35), and with one previous study of adult BED noting hypoconnectivity between reward and cognitive control networks (22), we examined rsFC within and between reward and cognitive control networks in pre-adolescent children with BED. Contrary to hypotheses, our findings suggest no alterations in functional connectivity *within* reward and cognitive control networks, respectively. However, findings illustrated a diffuse dysconnectivity *between* nodes of the reward and cognitive control networks in pre-adolescent children with BED, indicating for the first time that abnormalities in the functional synergy between reward and cognitive control networks may be evident in children with BED as young as nine years of age.

Specifically, we noted reduced connectivity between the dlPFC and amygdala, and between the anterior cingulate cortex and orbitofrontal cortex in pre-adolescent children with BED. These findings illustrating hypoconnectivity between reward and cognitive control networks accord with theoretical models of the psychopathology of BED, which posit a critical interplay between reward and impulse control-related processes (35). Essentially, it is thought that the increased food intake characteristic of BED typically develops when an increased reward value is placed on food, therefore facilitating a higher drive to eat, and persists over time if this heightened drive cannot be inhibited (10). Our findings replicate recent findings in (i) adults with BED, noting hypoconnectivity between seed regions involved in reward processing, such as the NAcc and dorsal caudate, and prefrontal regions involved in cognitive control, such as the superior frontal gyrus (22) and (ii) pre-adolescent children who overeat, noting that eating in the absence of hunger is associated with the functional connectivity between the caudate and dlPFC (36). Crucially, our hypothesis-driven findings extend these earlier findings by implicating additional regions of dysconnectivity between reward and cognitive control regions – namely the (i) dlPFC and amygdala, and (ii) dACC and OFC.

These hypotheses-driven seed-to-seed findings are further supported by our exploratory seed-to-voxel whole-brain analyses. These statistically significant and corrected findings suggest broad dysconnectivity between reward and cognitive control circuits, illustrating that the dysconnectivity from (i) the amygdala to regions in the dlPFC and SFG, (ii) the OFC to regions in the dlPFC, ACC, vmPFC and PCC (iii) the ACC to regions in the vmPFC, SFG, and OFC, and (iv) from the dlPFC to regions in the OFC and vmPFC, rank among the most dysconnected of any voxels in the whole brain from these respective seed regions. Cumulatively, our findings suggest that broad dysconnectivity between reward and cognitive control networks may characterize BED, which may have profound implications for the adaptive responding to rewarding cues, which rests on an intricate calibration of approach behaviors and cognitive/behavioral inhibition. The broadly reduced synchronicity of these networks in pre-adolescent children with early-onset BED, and the notion that this dysconnectivity is associated with binge frequency and volume in adult BED (22, 36), suggests that these processes and networks are particularly important targets for treatment efforts among children with BED.

With respect to our assessment of the functional connectivity of nodes *within* the reward network, we found no evidence of dysconnectivity in our sample of pre-adolescent children with BED, relative to weight-matched controls. This accords with previous whole-brain seed-to-voxel analyses in adults with BED, which revealed no altered connectivity patterns within nodes of the reward network, relative to weight-matched controls (22). Interestingly, previous studies of functional connectivity in those reporting binge type behaviors in obese versus non-obese populations have revealed mixed findings with regard to within reward network functional connectivity. For instance, in a study comparing obese individuals (some of whom had BED) and non-obese individuals, *reduced* connectivity was noted in obese participants between nodes within the basal ganglia reward circuit, including the pallidum, putamen and amygdala, which was negatively correlated with patient weight and binge eating symptoms (23). In contrast, food addiction among those with obesity has been associated with *increased* connectivity between nodes within the reward network, relative to obese individuals without food addiction (37), and was associated with both the frequency and intensity of general food cravings in obese populations (37). Further, functional connectivity analyses in non-obese adults and children who report binge and overeating (36) episodes, respectively, have revealed non-disturbed functional connectivity within the reward network (37), raising the interesting possibility that reward network functional abnormalities may be gated by bodyweight among those who engage in binge type behaviors. Standalone alterations in functional connectivity of the reward network have been well demonstrated among those with obesity, irrespective of the presence of binge eating (37), and recent machine learning efforts have outlined a robust obesity functional connectivity phenotype involving hyperconnectivity within the reward network, which correlates with body mass, waist circumference, and waist-to-hip ratio. Since controls in the present study were BMI-matched, it is noteworthy that reward network functional connectivity in overweight pre-adolescent children with BED was not discrepant from that of overweight pre-adolescent children without BED.

With respect to connectivity *within* cognitive control networks, we found no evidence of dysconnectivity in BED. This echoes findings from previous studies of BED (22, 23), and several studies assessing functional connectivity and overeating in obese and non-obese populations. Notably, one previous study of restrained eaters noted diminished inter-hemispheric dlPFC functional connectivity, relative to non-restrained eaters, which was associated with greater binge-type eating symptoms (38). While the present study did not assess inter-hemispheric functional connectivity, our findings suggesting that inter-node connectivity within cognitive control networks is not fundamentally disturbed in those with BED, relative to weight-matched controls. This accords with studies demonstrating comparable performance in neuropsychological tests of inhibitory control among obese women with and without BED (11). However, even in the context of comparable task performance on measures of inhibitory control, some studies have noted distinct neural activity in prefrontal regions among obese individuals with BED, relative to obese individuals without BED, and non-obese controls (21), suggesting divergent neural substrates underpinning inhibitory control. Our findings suggest that any functional atypicality in neural activity among those with BED during tasks of inhibitory control do not stem from alterations in functional connectivity within the cognitive control network. However, with studies illustrating reliable differences in cognitive control between obese and non-obese samples (39), it is noteworthy that both cohorts in the present study were obese. A crucial next step lies in parsing functional differences among obese and non-obese BED.

Strengths of the present study include the relatively large sample size, which is among the largest neuroimaging studies of pediatric BED to date. Additionally, the focus on pre-adolescent BED offers important insights on the mechanistic underpinnings of early onset BED, which has been seldom studied. With much evidence noting the benefits of early intervention for ED (40), an explication of the precise mechanisms underpinning pediatric BED may help mitigate the suite of deleterious outcomes associated with its longer-term course (3, 4). Relatedly, and owing to the centrality of impulse control in theoretical models of BED (35), our assessment of pre-adolescent children accounted for developmental maturation, which is critical given the potentially discrepant development of the late-maturing prefrontal cortex. Lastly, diagnoses in this instance were based on the reports of parents and caregivers, rather than the self-reported accounts of 9-10-year-old children themselves. With a diagnosis of BED resting on an understanding of abstract cognitive constructs such as what constitutes a loss of control, and what is an objectively large amount of food relative to same-age peers, and with parents being reliable informants of child and adolescent ED psychopathology (29), parent diagnostic reports were adopted. Limitations of the present study are also noteworthy. First, and owing to the nature of the parent ABCD dataset, no dimensional measure of ED symptomatology was included. It is critical that future studies investigate how these patterns of functional dysconnectivity relate to ED symptoms among those with BED. Secondly, food consumption prior to scanning was unknown. With evidence suggesting that neural activity in those with ED, and particularly in reward-related regions, may differ during sated and hungry states (14, 21, 36), the inability to control for pre-scan food consumption may impact the generalizability of the findings. Lastly, the variability in BMI among those with BED in our sample was wide, whereas the variability among those in our control sample was narrow. This means that a potentially disparate sample (in terms of BMI) of individuals with BED were compared to a highly circumscribed sample of control individuals. Further research may seek to parse the effect of patient weight upon functional connectivity patterns among those demonstrating BED and binge eating behaviors, given the variable findings among obese and non-obese individuals (23, 36, 37). Notwithstanding, these findings offer novel insights into the neural architecture of early-onset BED, and suggest that rather than inherent functional connectivity abnormalities within reward and cognitive control networks, early-onset BED may be characterized by abnormalities in the functional synchrony between reward and cognitive control networks.

## Additional Information

The ABCD Study was supported by the National Institutes of Health and additional federal partners under award numbers U01DA041022, U01DA041025, U01DA041028, U01DA041048, U01DA041089, U01DA041093, U01DA041106, U01DA041117, U01DA041120, U01DA041134, U01DA041148, U01DA041156, U01DA041174, U24DA041123, and U24DA041147. A full list of supporters is available at https://abcdstudy.org/nihcollaborators. A listing of participating sites and a complete listing of the study investigators can be found at https://abcdstudy.org/principal-investigators.html. ABCD consortium investigators designed and implemented the study and/or provided data but did not necessarily participate in the analysis or writing of this report.

